# Amot regulates neuronal dendritic tree through Yap1

**DOI:** 10.1101/345264

**Authors:** Katarzyna O. Rojek, Joanna Krzemień, Hubert Doleżyczek, Paweł M. Boguszewski, Leszek Kaczmarek, Witold Konopka, Marcin Rylski, Jacek Jaworski, Lars Holmgren, Tomasz J. Prószyński

## Abstract

The Amot-Yap1 complex plays a major role in the regulation of cell contact inhibition, cellular polarity and growth. However, the function of Angiomotin (Amot) and Hippo pathway transcription co-activator Yap1 in the central nervous system remains unclear. In this study, we demonstrate that Amot is a critical mediator of dendritic morphogenesis in cultured hippocampal cells and Purkinje cells in the brain. Amot function in developing hippocampal neurons depends on interactions with Yap1, which is also indispensable for dendrite growth and arborization in vitro. Conditional deletion of Amot or Yap1 in neurons leads to impaired morphogenesis of Purkinje cell dendritic trees, decreased cerebellar size, and causes defects in locomotor coordination of mutant animals. Thus, our studies identified Amot and Yap1 as novel regulators of dendritic tree morphogenesis.

## INTRODUCTION

Neurons are highly polarized cells with specialized axonal and somatodendritic compartments. The architecture of dendritic arbors at which information from other neurons is received is crucial for the integration and processing of signals (London and Hausser, 2005; Segev and London, 2000). Although the characteristic dendritic pattern is driven by both intrinsic mechanisms and external signals (Bartlett and Banker, 1984a; Bartlett and Banker, 1984b; Gray et al., 2004), the molecular processes that underlie the formation of dendritic trees have been studied mostly in cultured hippocampal neurons under conditions in which external signals are limited. Abnormalities in dendritic arborization are associated with numerous neurological disorders, such as schizophrenia, epilepsy, Alzheimer’s disease, and autism spectrum disorders (Bernardinelli et al., 2014; Bourgeron, 2015; Li et al., 2015; Tampellini, 2015; Wu et al., 2015). Thus, a better understanding of the molecular processes that regulate dendritic tree patterning could facilitate the development of new treatment modalities for various neurological and psychiatric conditions.

Angiomotin (Amot) protein together with the closely related Angiomotin-like 1 (Amotl1) and Angiomotin-like 2 (Amotl2) constitute a family of scaffold proteins called Angiomotins or Motins (Bratt et al., 2002). Amot is the most characterized member of the Angiomotin family. This protein was shown to regulate many physiological and pathological processes including angiogenesis, cell polarity and migration, as well as cancer cell progression (Aase et al., 2007; Ernkvist et al., 2009; Hsu et al., 2015; Lv et al., 2016; Lv et al., 2015; Troyanovsky et al., 2001; Wells et al., 2006). However, one of the most characterized functions of Amot and other Motins is regulation of the Hippo signaling pathway.

The Hippo signaling pathway controls cell growth and differentiation (Gumbiner and Kim, 2014; Yu et al., 2015; Zhao et al., 2010; Zhao et al., 2007). Core to this pathway is a cascade of kinases, Mst and Lats, regulating phosphorylation of the transcription co-activators Yap1 and Taz, two major downstream effectors of the Hippo pathway (Callus et al., 2006; Chan et al., 2005; Hao et al., 2008; Lei et al., 2008; Zhao et al., 2010; Zhao et al., 2007). Phosphorylation deactivates Yap1 by inhibiting its translocation to the nucleus (Zhao et al., 2010; Zhao et al., 2007) where it interacts with TEA domain (TEAD) transcription factors to induce the expression of Hippo pathway-dependent genes (Vassilev et al., 2001; Zhao et al., 2010; Zhao et al., 2008). Amot has been shown to directly and strongly interact with Yap1 (Zhao et al., 2011). The function of Amot in the regulation of Yap1 appears to be both tissue- and cell type-specific (Chan et al., 2011; Lv et al., 2015; Yi et al., 2013; Zhao et al., 2011). For instance, in MDCK epithelial cells and human embryonic kidney 293 (HEK-293) cells, Amot inhibits Yap1-dependent transcription. In contrast, in hepatocarcinoma and breast cancer cells, Amot induces the transcription of TEAD-target genes (Chan et al., 2011; Lv et al., 2015; Yi et al., 2013; Zhao et al., 2011). Both Amot and Yap1 have been shown to regulate contact-mediated inhibition of cell proliferation and control organ size and growth (Gumbiner and Kim, 2014; Lv et al., 2016; Zhao et al., 2011; Zhao et al., 2007).

The function of Amot protein in neurons is poorly understood. Recent studies demonstrated that Amot is localized to dendritic spines in cultured hippocampal neurons where it controls the integrity of postsynaptic density by regulating MUPP1 and PSD-95 (Wigerius et al., 2018). Schanzenbächer and colleagues (Schanzenbacher et al., 2016) have reported that Amot is associated with neuronal homeostatic scaling and autism spectrum disorders. In the present study, we report that Amot is enriched at synapses of mature neurons; however, at earlier developmental stages Amot is localized to axonal and dendritic processes and is required for the proper growth and development of dendritic arbors both *in vitro* and *in vivo*. We found that Amot in neurons interacts with Yap1, which similarly to Amot, localizes to neurites in young neurons and translocates to synapses in mature cells. Additionally, we found that Yap1 is critical for dendritic tree growth and arborization. Thus, our study unravels novel molecular machinery regulating neuronal dendritic tree organization.

## RESULTS

### Amot expression and localization in neurons

Amot has been previously shown to be ubiquitously expressed in various rat brain structures and to be localized to synapses in cultured mature neurons (Wigerius et al., 2018). Similarly, our Western blotting analysis of brain homogenates revealed high levels of Amot expression in different regions of mouse brain (Fig. 1A). The full-length isoform of Amot (p130) was predominantly expressed in the brain but the shorter isoform (p80) lacking the N’-terminal domain was also produced in lower amounts. To study Amot localization in the brain we performed immunohistochemical analysis of cryostat brain sections. These experiment revealed that Amot concentrates at synapses in the CA1 hippocampal region (Fig. 1B) and Purkinje cells in cerebellum (Fig. 1C), where its immunoreactivity overlaps with anti-synaptophysin staining.

**Figure 1.**
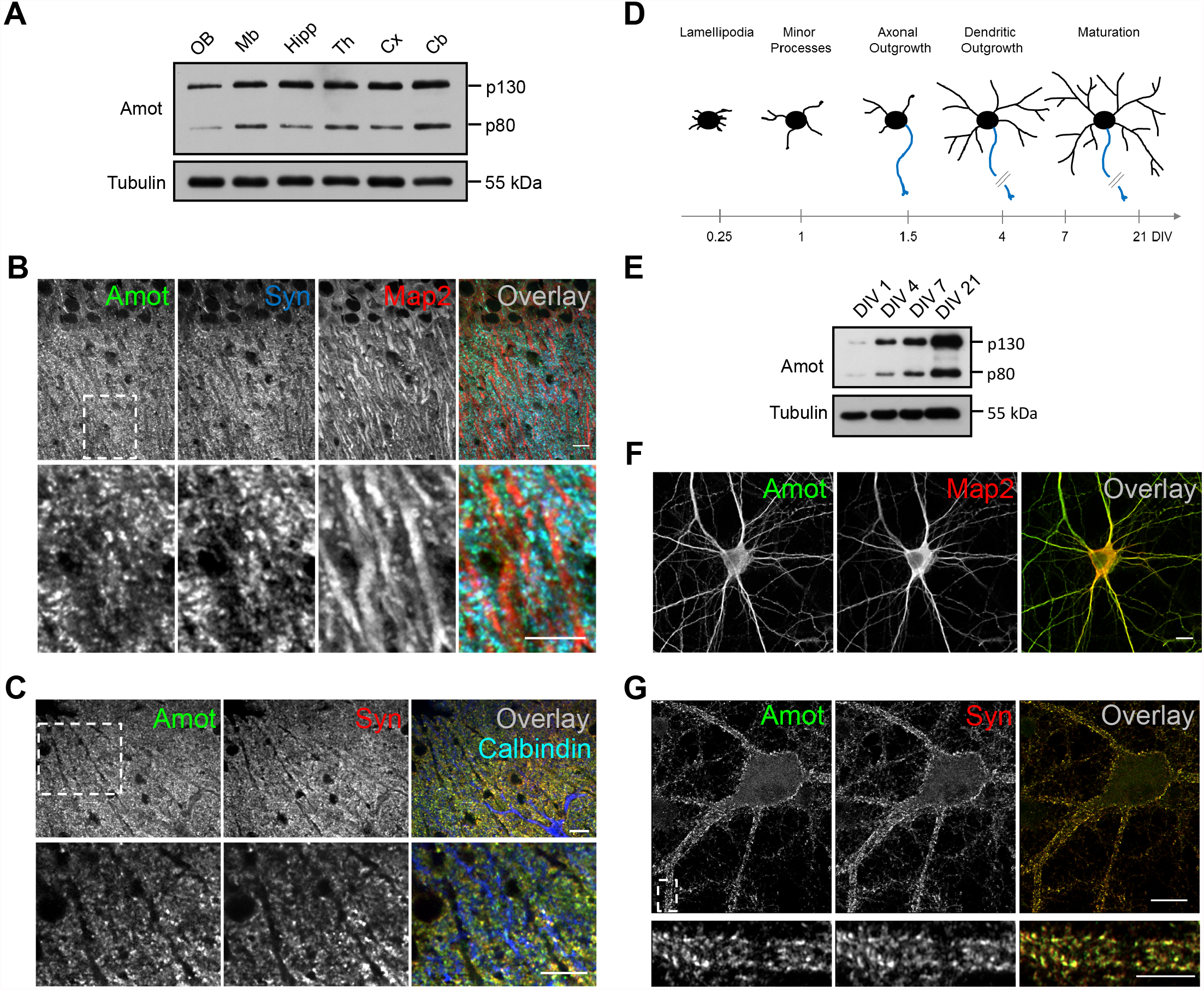
Expression of Angiomotin proteins in neurons. (**A**) Western blot analysis of Amot expression in different regions of the mouse brain. OB, olfactory bulbs; Mb, midbrain; Hipp, hippocampus; Th, thalamus; Cx, cortex; Cb, cerebellum. (**B**, **C**) Amot localization in neurons in the cryostat sections of P30 mouse brain. (**B**) CA1 region of the hippocampus stained for Amot (green), Synaptophysin (Syn, blue), and Map2 (red). (**C**) Purkinje cells dendritic tree in cerebellum stained for Amot (green), Synaptophysin (Syn, red), and Calbindin (blue). Lower panels show higher magnification of the boxed areas in the upper panels. (**D**) Model of hippocampal neuron development *in vitro*. Development of somatodendritic compartment (black) and axon (blue) is correlated with days *in vitro* (DIVs) on the axis at the bottom. (**E**) Amot expression increases at later stages of neuronal development. Western blot analysis of neuronal extracts collected at the indicated DIVs. (**F**, **G**) Localization of Amot protein in cultured neurons. (**F**) Rat hippocampal neurons (DIV 7) were immunolabeled for Amot (green) and Map2 (red).(**G**) Amot localization in DIV 21 cultured rat hippocampal neurons; Amot (green) and Synaptophysin (Syn, red). The lower panel shows higher magnification of boxed area in the upper panel. Scale bar = 10 μm on **B**, **C**, **F**, and **G** (upper panel) and 5 μm on **G** (lower panel).

We next studied in details expression and localization of Amot in hippocampal neurons cultured *in vitro*. Western blot analysis revealed that Amot expression gradually increased during neuronal differentiation (Fig. 1D, E) and was particularly high at the time of intensive dendritogenesis on day four to seven *in vitro* (DIV 4-7). The increase of expression was similar for both isoforms of Amot p130 and p80. Our immunocytochemical analysis of cultured rat hippocampal neurons on DIV 7 showed that at this developmental stage Amot was mostly concentrated in dendrites and axons (Fig. 1F). Similar localization to dendrites was observed in DIV 7 hippocampal neurons overexpressing GFP-tagged version of Amot (Fig. S1). At later stages of neuronal development (DIV 21) Amot re-localized to synaptic compartments, where it colocalized with synaptophysin puncta, as revealed by Airyscan microscopy (Fig. 1G).

### Amot regulates dendritic tree development in cultured hippocampal neurons

Amot’s localization within dendritic and axonal compartments during neuronal maturation and its higher expression at the time of neurite outgrowth suggest that it could regulate the development of neuronal processes. To investigate this possibility, we performed RNA interference (RNAi)-mediated silencing of Amot expression in DIV7 cultured hippocampal neurons. We cloned short-hairpin RNA (shRNA) plasmids and selected the one with the highest knockdown efficiency in neurons, which was confirmed by quantitative real-time polymerase chain reaction (qRT-PCR) and Western blot (Fig. 2A, B). To visualize the morphology of individual neurons, we co-transfected cells with the shRNA and a plasmid expressing monomeric red fluorescent protein (mRFP) under the control of β-actin regulatory elements. As expected, neurons that were transfected with Amot shRNA plasmid had severe defects in dendritic tree arborization compared to control cells that were transfected with an empty vector (Fig. 2C). These cells had a shorter total dendrite length (TDL; sum of all dendrite lengths) by approximately 75% compared with control neurons (Fig. 2D) and formed significantly less complex dendritic arbors, verified by Sholl analysis (Fig. 2E). Sholl analysis indicates the number of dendrites that cross circles at various radial distances from the cell soma and describes dendritic arbor complexity and the area of a dendritic field (Sholl, 1953). The observed phenotypes of neuron morphology were specific to Amot rather than an off-target effect of overexpressed shRNA, because the phenotype could be reversed (i.e., rescued) by the ectopic expression of a mouse Amot construct lacking nucleotide sequence targeted by shRNA (Fig. 2C-E). The overexpression of Amot alone had no significant effects on dendritic tree development (Fig. 2D, E).

**Figure 2.**
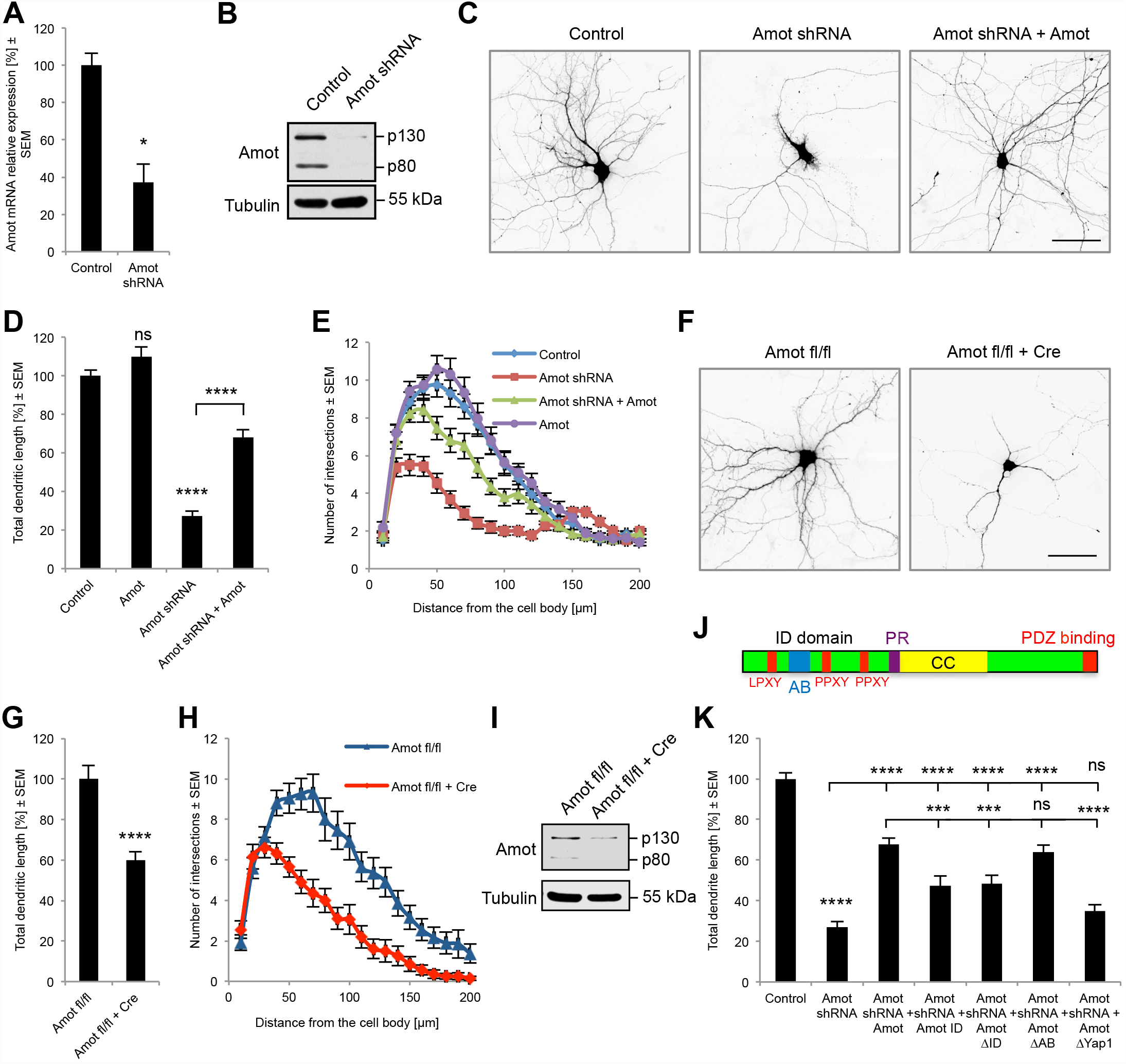
Amot knockdown in cultured hippocampal neurons affects dendritic tree organization. Validation of Amot knockdown efficiency in neurons by qRT-PCR (n = 3 per group; P = 0.0191) (**A**) and Western blot (**B**). (**C**) Rat hippocampal neurons transfected on DIV 7 with an empty pLKO plasmid (control), a plasmid with Amot shRNA, and a plasmid with Amot shRNA and a plasmid expressing mouse Amot-GFP (Amot shRNA + Amot; rescue experiment). Quantification of TDL shown as percentage of Control (**D**) and Sholl analysis (**E**) of hippocampal neurons transfected with plasmids as in (**C**) or transfected with Amot-GFP plasmid. Control *n* = 60, Amot *n* = 38, Amot shRNA *n* = 60, Amot shRNA + Amot *n* = 58. To Control; P = 0.0820, P < 0.0001; to Amot shRNA; P < 0.0001. (**F**) Representative images of hippocampal neurons obtained from Amot fl/fl mice transfected with a control vector or a plasmid that expressed Cre. Quantification of TDL shown as percentage of control (**G**) and Sholl analysis (**H**) of dendritic trees of hippocampal Amot fl/fl neurons transfected with either a control vector (*n* = 33) or a Cre-expressing plasmid (*n* = 40); P < 0.0001. (**I**) Cre expression in Amot fl/fl hippocampal neurons led to the downregulation of Amot expression in the Western blot analysis. (**J**) Domain architecture of Amot protein. Amot ID domain contains LPXY and PPXY motifs (red) that bind Yap1 and the actin-binding motif (AB, blue). ID domain, intrinsically disordered domain; CC, coiled-coil domain, PR, prolin rich region. (**K**) Quantification of TDL of Amot knockdown rat hippocampal neurons transfected with indicated Amot constructs; values shown as percentage of Control. Control cells were transfected with an empty pLKO vector. Control *n* = 107, Amot shRNA *n* = 80, Amot shRNA + Amot *n* = 100, Amot shRNA + Amot ID domain *n* = 37, Amot shRNA + Amot ΔID domain *n* = 45, Amot shRNA + Amot ΔAB *n* = 59, Amot shRNA + Amot ΔYap1 *n* = 59. To control; P < 0.0001; to Amot shRNA; P < 0.0001, P < 0.0001, P < 0.0001, P < 0.0001, P = 0.0554; to Amot shRNA + Amot; P = 0.0009, P = 0.0006, P = 0.4416, P < 0.0001. For morphological analysis, neurons were transfected with a plasmid expressing mRFP to visualize cell morphology and a GFP plasmid (if needed) to maintain plasmid ratio. Images were obtained at least from three independent cultures. Statistical significance was analyzed using two-tailed and unpaired *t*-test (**A**, **D**, **G**, **K**) and two-way analysis of variance followed by Bonferroni’s post hoc test (**E**, **H**). See Supplementary Table S1 for detailed statistics for **E** and **H**. ns, not significant; \**P* < 0.05, \*\*\**P* < 0.001, \*\*\*\**P* < 0.0001. Bars represent mean ± standard error of the mean (SEM). Scale bars = 50 μm.

To independently confirm the involvement of Amot in dendritic tree outgrowth, we examined the morphology of cultured hippocampal neurons that were obtained from conditional Amot knockout mice (Amot flox/flox; see also Fig. 5 and the description in the following paragraphs) that were co-transfected with a GFP-encoding plasmid to visualize their morphology and either a Cre recombinase-expressing plasmid or control plasmid without Cre (Fig. 2F). Neurons that were transfected with the Cre plasmid (Fig. S2A) had a shorter TDL by approximately 60% and simplified dendritic tree morphology as revealed by Sholl analysis (Fig. 2G, H). To confirm the reduction of Amot expression, we performed Western blot analysis of extracts from Amot flox/flox cortical neurons that were nucleofected with a plasmid that encoded Cre recombinase or a control vector. We found that Cre expression in Amot flox/flox neurons reduced Amot protein levels (Fig 2I), although some residual expression was still detected, likely because of the ∼50% transfection efficiency in the nucleofection experiments. Cre expression in cultured neurons that were obtained from wildtype mice did not decrease Amot expression and had no effect on dendrite outgrowth (Fig. S2B-D). Altogether, these results support the conclusion that Amot is required for proper dendritic arbor morphology in developing hippocampal neurons.

To identify the Amot protein region that is responsible for regulating dendritic outgrowth, we rescued the Amot knockdown phenotype by ectopically expressing truncated Amot constructs. The N-terminal intrinsic disordered (ID) domain of Amot (Fig. 2J) is known to interact directly with F-actin through a 34-amino-acid sequence (Mana-Capelli et al., 2014) and contains two PPXY motifs and a LPXY motif that bind the Hippo pathway transcription coactivator Yap1 (Yi et al., 2013; Zhao et al., 2011). The transfection of Amot-depleted neurons with constructs that encoded either the ID domain alone (Amot ID) or Amot with a deleted ID domain (Amot ΔID) led to the partial restoration of dendritic tree arborization compared with the expression of full-length Amot (Fig. 2K). This observation suggests that Amot ID domain plays an important role in the regulation of dendritic processes, but other domains could also be involved. We then investigated whether the Amot-regulated organization of neuronal dendritic trees depends on the actin-binding site (Ernkvist et al., 2006; Mana-Capelli et al., 2014) that is centrally located within the ID domain (see Fig. 2J). The expression of Amot mutant protein with a deleted actin-binding region (Amot ΔAB) rescued the phenotype of Amot depletion to an extent that was comparable to the expression of full-length Amot protein (Fig. 2K). To study the importance of the Yap1-binding motifs located in the Amot ID domain, we expressed Amot mutant with impaired binding to Yap1 in Amot knockdown neurons. The expression of Amot defective in Yap1 binding (Amot ΔYap1) failed to rescue impairments of dendritic processes outgrowth observed in Amot depleted neurons. Importantly, all of the Amot constructs that were used in the phenotype rescue experiments were expressed in neurons at similar levels (Fig. S3A, B). These results suggest that the interaction with Yap1 plays an important role in Amot-dependent regulation of dendritic tree.

### Expression and localization of Yap1 in neurons

Although Yap1 has been previously shown to be expressed in progenitor cells in the brain where it regulates proliferation, migration, and differentiation under both physiological and pathological conditions, the function of Yap1 has not yet been characterized in vertebrate neurons (Huang et al., 2016a; Huang et al., 2016b; Mao et al., 2016; Yamanishi et al., 2017). Our Western blot analysis revealed that Yap1 is widely expressed in different brain structures with the highest level detected in cerebellum (Fig. 3A). To verify the interaction between Amot and Yap1 we precipitated Yap1 from mouse brain extract and observed that Amot co-precipitates with Yap1 but not with control beads (Fig 3B), what suggests an interaction between these two proteins. This is in agreement with observations from several other groups that have demonstrated direct interaction between Amot and Yap1 in different cell types (Chan et al., 2011; Yi et al., 2013; Zhao et al., 2011). To study localization of Yap1 we immunohistochemically analyzed cryostat sections of the mouse P30 hippocampus (Fig. 3C) and cerebellum (Fig. 3D). Yap1 immunoreactivity was observed predominantly at the synaptic compartments where it colocalized with synaptophysin puncta.

**Figure 3.**
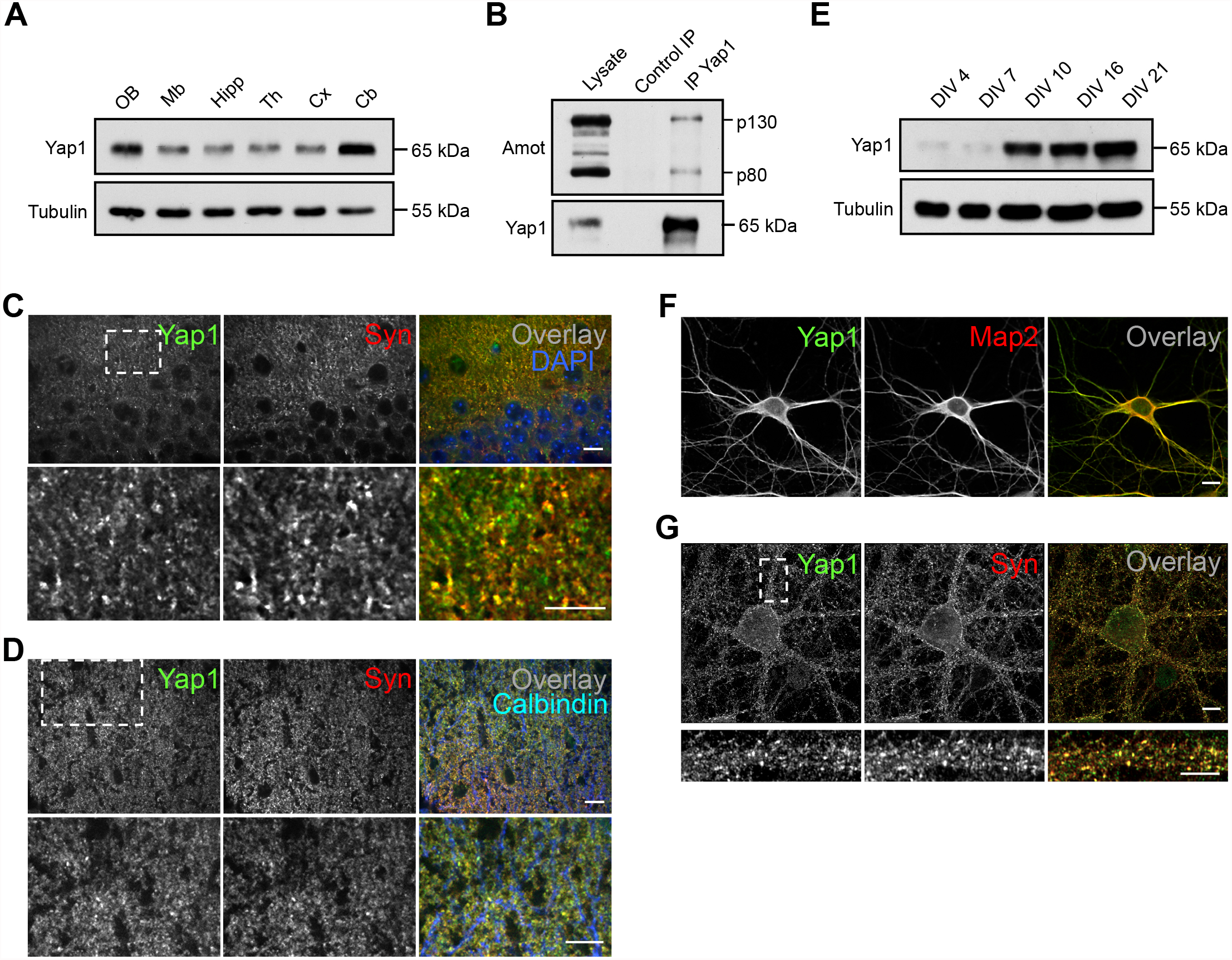
Expression and localization of Yap1 in neurons. (**A**) Western blot analysis of Yap1 expression in different regions of the mouse brain. OB, olfactory bulbs; Mb, midbrain; Hipp, hippocampus; Th, thalamus; Cx, cortex; Cb, cerebellum. (**B**) Analysis of Amot interaction with Yap1 through co-immunoprecipitation. Magnetic beads coated with anti-Yap1 antibody or uncoated (control beads) were used to precipitate Yap1 from mouse brain homogenate. Precipitates were analyzed for Amot and Yap1 by Western blot. (**C**, **D**) Yap1 localization in the brain. Cryostat sections of P30 mouse hippocampus (**C**) and cerebellum (**D**) stained for Yap1 (green) and Synaptophysin (Syn, red). Overlay images additionally show nuclei visualized with DAPI (blue on **C**) and Purkinje cells dendritic processes visualized with anti-Calbindin antibody (blue on **D**). Lower panels on **C** and **D** show higher magnification of the boxed areas in the upper panel. (**E**) Yap1 expression increases at later stages of neuronal development. Western blot analysis of neuronal extracts collected at the indicated DIVs. (**F**) Yap1 localizes to dendritic processes in young cultured neuronal cells. Rat hippocampal neurons (DIV 8) were immunolabeled for Yap1 (green) and Map2 (red). (**G**) Yap1 localizes to synapses in mature cultured neuronal cells. Rat hippocampal neurons (DIV 21) were immunolabeled for Yap1 (green) and Synaptophysin (Syn, red). The lower panel shows higher magnification of boxed area in the upper panel. Scale bar = 10 μm on **C**, **D**, **F**, and **G** (upper panel) and 5 μm on **G** (lower panel).

We next studied expression and localization of Yap1 in hippocampal neurons cultured *in vitro*. Western blot analysis revealed that Yap1 expression is increased after DIV 7 (Fig. 3E). Immunocytochemical analysis of cultured rat hippocampal neurons on DIV 8 showed that at this developmental stage Yap1 was mostly concentrated in dendrites and axons (Fig. 3F). At later stages of neuronal development (DIV 21) Yap1 re-localized to synaptic compartments, where it colocalized with synaptophysin puncta, as revealed by Airyscan microscopy (Fig. 3G). Thus, Yap1 localization resembles Amot at the early and late stages of neuronal development.

### Yap1 regulates dendritic tree organization

Since Yap1 localized to dendrites in developing hippocampal neurons and the ectopic expression of Amot protein with mutated Yap1 binding sites failed to rescue phenotypes observed in Amot depleted cells, we hypothesized that Amot may organize dendritic trees through Yap1.

To determine whether Yap1 plays a role in dendritic tree arbor development, we knocked down Yap1 expression in cultured rat hippocampal neurons by transfecting cells with shRNA, which has the highest knockdown efficiency in neurons (Fig. 4A) and examined their morphology. Hippocampal neurons transfected with a plasmid that encoded shRNA targeting rat Yap1 exhibited significant impairments in dendritic arbors, reflected by a 40% decrease in the TDL and a shift in the Sholl plots compared with control cells that were transfected with an empty vector (Fig. 4B-D). The negative effects of Yap1 knockdown on dendritic arbors were almost fully reversed by the ectopic expression of human Yap1 cDNA, which lacks the nucleotide sequence targeted by used shRNA designed against rat ortholog (Fig. 4B-D). One of the well-documented functions of Yap1 is regulation of gene expression. This depends on the interaction of Yap1 with TEA domain (TEAD) transcription factors, which upon interaction with co-activators bind to TEA sequences in the regulatory elements of many genes stimulating their transcription (Vassilev et al., 2001; Zhao et al., 2008). To check if the Yap1 function in the organization of dendritic processes involves interaction with TEAD, we attempted to rescue phenotypes observed in Yap1 knockdown neurons by ectopic expression of mutant Yap1 protein (Yap1 del60-89) lacking the TEAD-binding domain (Shao et al., 2014). Interestingly, the expression of the Yap1 del60-89 mutant also rescued the defects in dendritic tree outgrowth to an extent that was comparable to wildtype Yap1 (Fig. 4C, D). In this experiment, both Yap1 constructs were expressed at similar levels (Fig. S3C). These observations suggest that Yap1 plays a role in dendritogenesis that is independent of interaction with TEAD.

**Figure 4.**
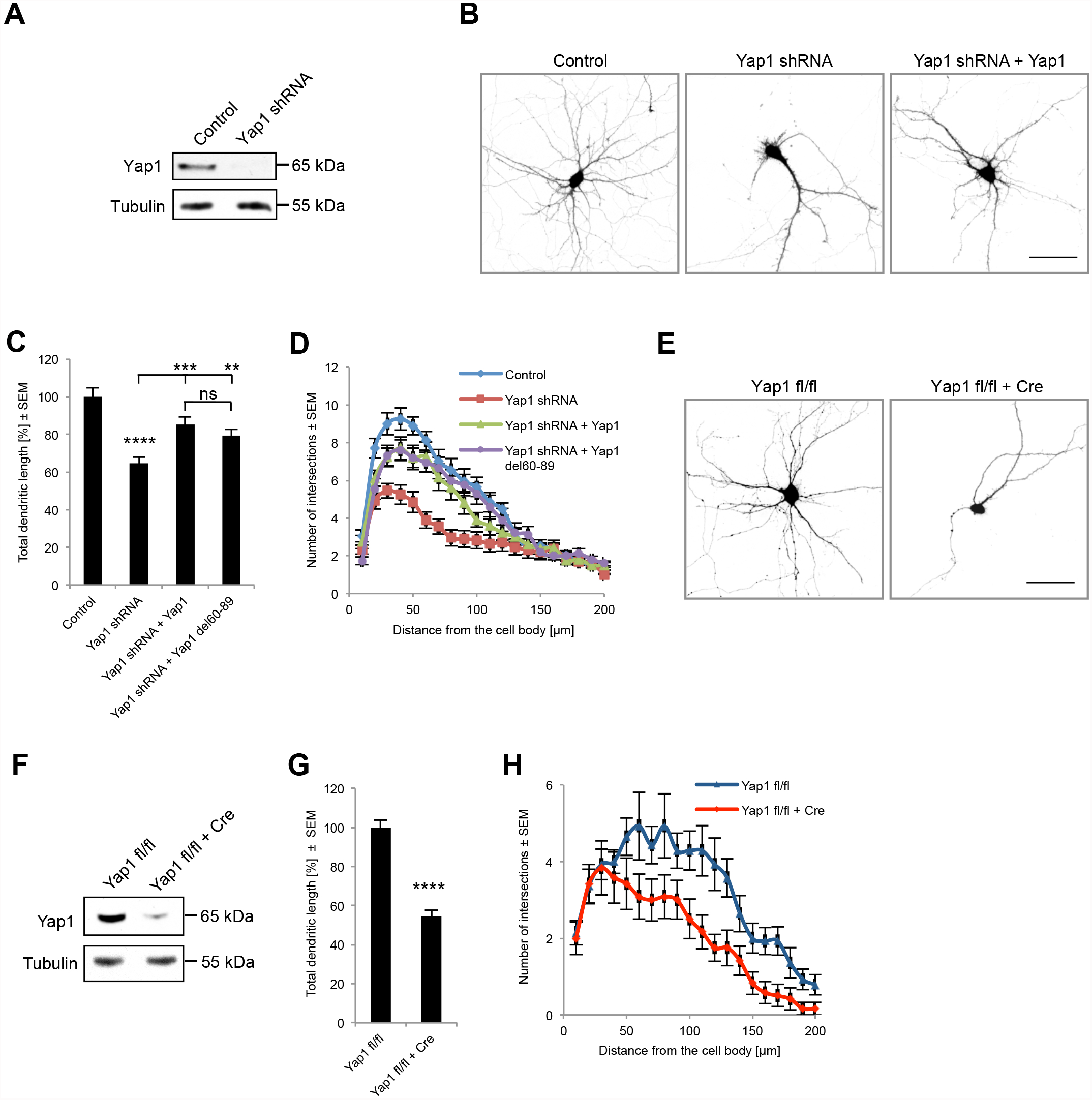
Yap1 regulates dendritic tree development. (**A**) Efficacy of Yap1 shRNA in neurons analyzed by Western blot. (**B**) Rat hippocampal neurons transfected on DIV 7 with an empty pLKO plasmid (control), a plasmid with Yap1 shRNA, and a plasmid with Yap1 shRNA and a plasmid expressing human Yap1-GFP (Yap1 shRNA + Yap1; rescue experiment). Quantification of TDL shown as percentage of Control (**C**) and Sholl analysis (**D**) of hippocampal neurons transfected with plasmids as in (**B**) or co-transfected with Yap1 shRNA and human Yap1-cDNA with a deleted TEAD-binding domain (Yap1 shRNA + Yap1 del60-89). Control *n* = 40, Yap1 shRNA *n* = 60, Yap1 shRNA + Yap1 *n* = 47, Yap1 shRNA + Yap1 del60-89 *n* = 51. To control; P < 0.0001; to Yap1 shRNA; P = 0.0002, P = 0.0034; to Yap1 shRNA + Yap1; P = 0.2715. (**E**) Representative images of hippocampal neurons obtained from Yap1 fl/fl mice transfected with a control vector or a plasmid that expressed Cre. (**F**) Cre expression in Yap1 fl/fl hippocampal neurons led to the downregulation of Yap1 expression in the Western blot analysis. Quantification of TDL shown as percentage of control (**G**) and Sholl analysis (**H**) of dendritic trees of hippocampal Yap1 fl/fl neurons transfected with either a control vector (*n* = 53) or a Cre-expressing plasmid (*n* = 53); P < 0.0001. For morphological analysis, neurons were transfected with a plasmid expressing mRFP to visualize cell morphology and a GFP plasmid (if needed) to maintain plasmid ratio. Images were obtained at least from three independent cultures. Statistical significance was analyzed using two-tailed unpaired *t*-test (**C**, **G**) and two-way analysis of variance followed by Bonferroni’s post hoc test (**D**, **H**). See Supplementary Table S1 for detailed statistics for **D** and **H**. ns, not significant; \*\**P* < 0.01, \*\*\**P* < 0.001, \*\*\*\**P* < 0.0001. Bars represent mean ± standard error of the mean (SEM). Scale bar = 50 μm

To independently confirm the role of Yap1 in dendritic tree growth, we examined the morphology of cultured hippocampal neurons that were obtained from Yap1 conditional knockout mice (Yap1 flox/flox; see also Fig. 7 and the description in the following paragraphs) and transfected with either a plasmid that expressed Cre recombinase or a control plasmid without Cre. We found that Cre expression in Yap1 flox/flox neurons decreased Yap1 expression, reduced the TDL by approximately 40%, and simplified dendritic tree morphology as revealed by Sholl analysis (Fig. 4E-H). In control neurons that were obtained from wildtype mice, Cre expression did not affect Yap1 protein levels or dendritic tree organization (Fig. S2C-E).

Altogether, our results suggest that Yap1 is expressed in neurons where it regulates dendritic tree outgrowth and arborization in a way that appears to be independent of interactions with the TEAD transcription factors.

### Characterization of Amot fl/fl;Syn-Cre mice

To investigate the role of Amot in the organization of neuronal cells *in vivo*, Amot fl/fl mice, in which exon 2 of Amot was flanked by loxP sites (Fig. 5A), were crossed with transgenic mice that expressed Cre recombinase under Synapsin 1 regulatory elements (Syn-Cre), leading to Cre expression that was restricted to neuronal cells. To confirm the activity and specificity of recombination in Syn-Cre mice, we crossed them with a STOP-tdTomato (STOP-Tom) reporter strain that contained the STOP-of-transcription sequence that was surrounded by LoxP sites and inserted between the CAG promoter and tdTomato coding sequence. STOP-Tom;Syn-Cre double transgenic mice exhibited strong tdTomato expression in the brain, with the highest fluorescence observed in the CA3 and dentate gyrus subregions of the hippocampus, midbrain, and cerebellum Purkinje cells but very low expression in the CA1 subregion of the hippocampus and cerebral cortex (Fig. 5B). To verify that Amot exon 2 was deleted by Cre recombinase in Amot fl/fl;Syn-Cre mice, we performed genotyping reactions with genomic DNA that was obtained from tail and brain. As shown in Fig. 5C, the electrophoresis of genotyping reactions revealed a 450 bp PCR fragment that corresponded to the deleted Amot sequence for DNA that was obtained from the brain of homozygous mice that expressed Syn-Cre (Fig. 5C). The Western blot analysis of hippocampal and cerebellar homogenates from homozygous mice with Syn-Cre transgene additionally confirmed reduced levels of Amot protein (Fig. 5D).

**Figure 5.**
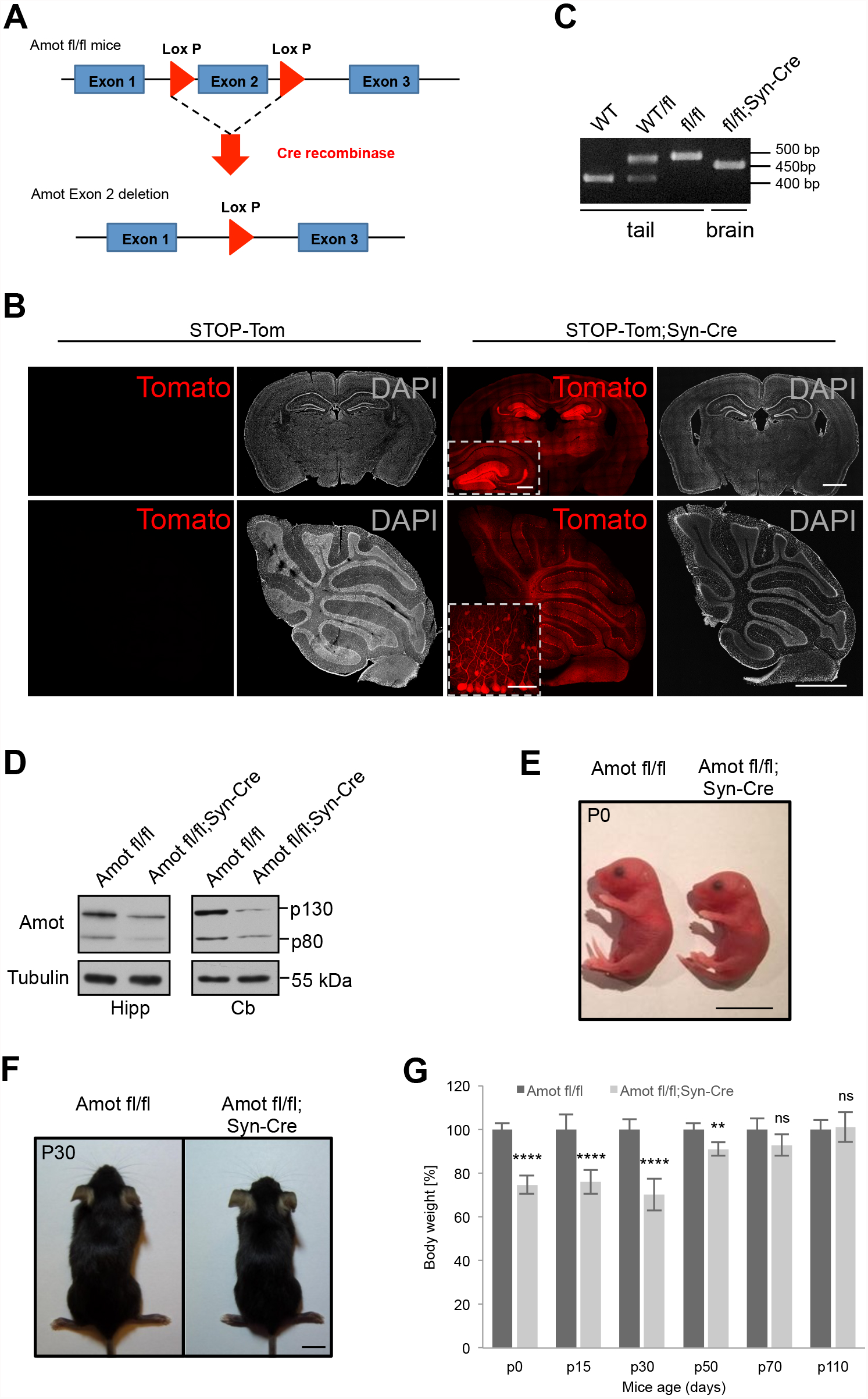
Mice with neuronal deletion of Amot. (**A**) Strategy for generation of Amot conditional knockout mice. The Amot flox allele (upper diagram) and recombined allele after Cre-mediated excision (lower diagram). (**B**) Cross-sections of brains from STOP-Tom;Syn-Cre or STOP-Tom (control) mice (P30) show high Cre activity in the hippocampus (upper panel) and cerebellum (lower panel). DAPI (gray) was used to visualize brain structure. Scale bars = 1 mm. Inserted squares show higher magnification images of hippocampus (upper panel, scale bar = 500 μm) and Purkinje cells (lower panel, scale bar = 50 μm). (**C**) Confirmation of Amot exon2 excision in the brain of Amot fl/fl;Syn-Cre mice assessed by PCR analysis of genomic DNA that was obtained from tail or brain. WT, wildtype control; WT/fl, Amot heterozygote; fl/fl, Amot homozygote mice. (**D**) Reduced Amot expression in the hippocampus and cerebellum of Amot fl/fl;Syn-Cre P30 mice analyzed by Western blot. (**E**, **F**) Representative images of Amot fl/fl;Syn-Cre and Amot fl/fl control littermates at the neonatal stage (**E**) or at P30 (**F**). P0, postnatal day 0; scale bars = 1 cm. (**G**) Weight analysis of Amot fl/fl and Amot fl/fl;Syn-Cre mice (*n* = 8, 9, 7, 3, 4, and 4 and *n* = 6, 4, 9, 6, 6, and 7, respectively) on the indicated days of development. Values shown as percentage of control (Amot fl/fl mice) weight at corresponding age. P < 0.0001, P < 0.0001, P < 0.0001, P = 0.0048, P = 0.0561 and P = 0.7799. Statistical significance was analyzed using two-tailed unpaired *t*-test. ns, not significant; \*\**P* < 0.01, \*\*\*\**P* < 0.0001. Bars represent mean ± standard deviation (SD).

Amot fl/fl;Syn-Cre mice were born at the expected Mendelian ratio and exhibited no signs of lethality. At birth, the pups of homozygous mice that expressed Cre, however, were significantly smaller and had a lower birth weight compared with their littermate controls of the same sex (Fig. 5E-G). These weight differences were apparent until approximately postnatal day 70 (P70). Afterward, Amot fl/fl;Syn-Cre mice grew to the same size and weight as Amot fl/fl mice, which were similar to wildtype animals (Fig. 5G).

### Neuronal deletion of Amot affects cerebellar morphology and the development of dendritic trees in Purkinje cells

We then analyzed the effects of Amot deletion in neurons on brain morphology. We made cryostat sections of brains that were collected from juvenile (P12) and early adult (P30) animals. Amot fl/fl;Syn-Cre mice did not exhibit apparent differences in size or gross morphology of the brain compared with littermate controls (Fig. 6A, B). However, the cerebellum appeared to be smaller (Fig. 6B) and had reduced weight (Fig. 6C) when compared with cerebellum of littermate controls. Because of this observation, and because the Syn-Cre activity was more homogenously distributed in the cerebellum and incomplete in the hippocampus (Fig. 5B), we focused our further experiments on the cerebellum.

**Figure 6.**
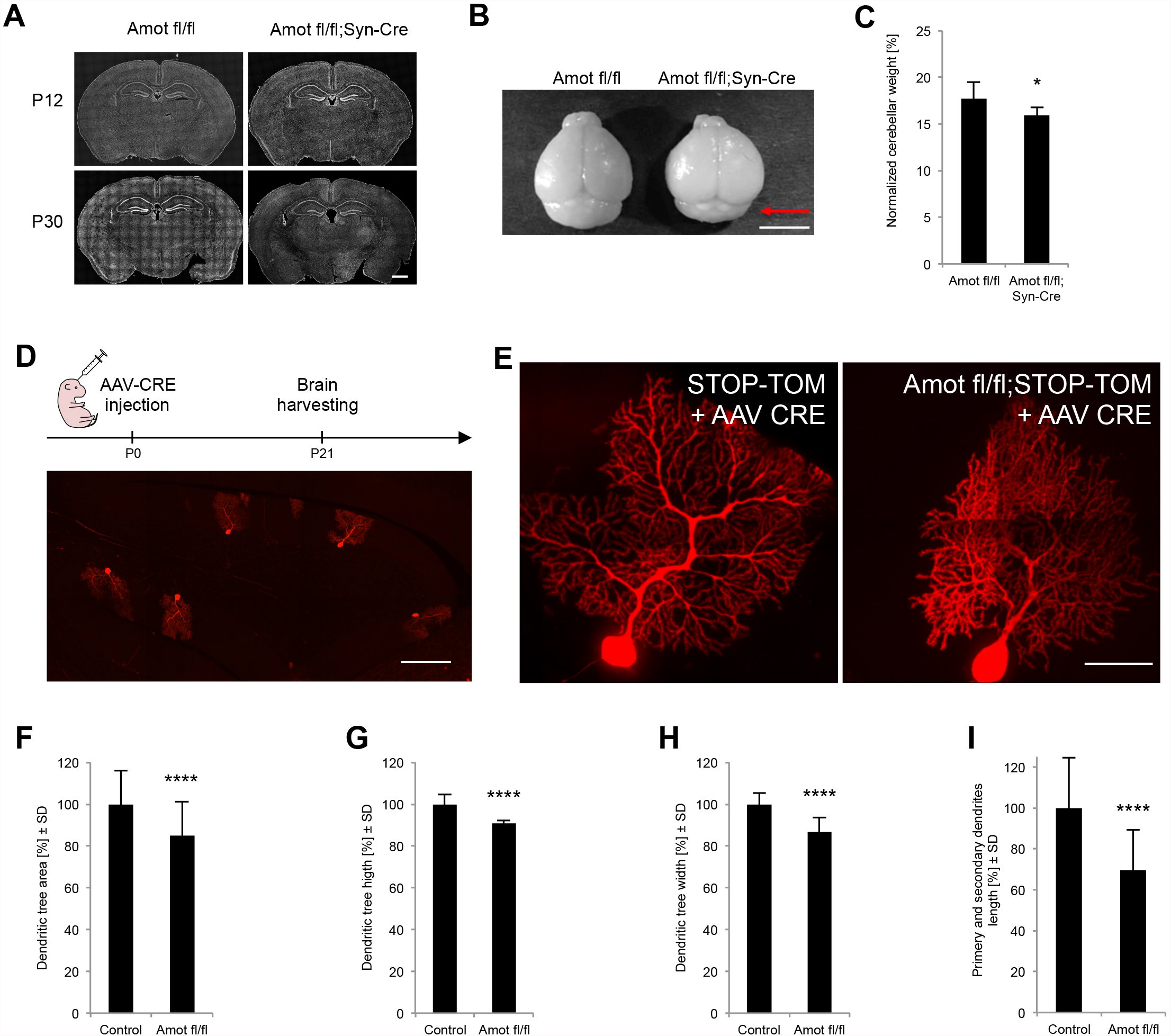
Amot deletion in neurons affects cerebellar size and impairs Purkinje cell dendritic tree morphology. (**A**) Coronal sections of Amot fl/fl and Amot fl/fl;Syn-Cre brains on P12 and P30. Scale bars = 1 mm. (**B**) Amot fl/fl;Syn-Cre mice exhibited a smaller cerebellum compared to control littermates (Amot fl/fl). Scale bars = 5 mm. (**C**) Quantitative analysis of the cerebellum weight of Amot fl/fl (*n* = 9) and Amot fl/fl;Syn-Cre (*n* = 6) mice; P = 0.0411; measurements were normalized to the whole brain weight. (**D**) Cre expression in single Purkinje cells in STOP-Tom reporter mice. Upper panel shows schematic illustration of the experimental procedure. P0 STOP-Tom pups were injected with AAV8-Cre virus and sacrificed on P21. The lower panel shows single Purkinje cells labeled in a sagittal section of the cerebellum from STOP-Tom mice after the AAV8-Cre injection. Scale bar = 200 μm. (**E**) Representative images of Purkinje cells from STOP-Tom and Amot fl/fl;STOP-Tom mice infected with AAV8-Cre. Scale bar = 50 μm. Quantitative analysis of dendritic tree area (**F**), height (**G**), width (**H**), and primary and secondary dendrites length (**I**) of Purkinje cells from STOP-Tom (*n* = 45 cells and 4 mice) and Amot fl/fl;STOP-Tom (*n* = 43 cells and 4 mice) brains infected with AAV8-Cre; values shown as percentage of Control. P < 0.0001, P < 0.0001, P < 0.0001 and P < 0.0001. Statistical significance was analyzed using two-tailed unpaired *t*-tests. \**P* < 0.05, \*\*\*\**P* < 0.0001. Bars represent mean ± standard deviation (SD).

To enable visualization of the morphology of individual Amot^−/−^ Purkinje cells, we intracerebrally injected newborn Amot fl/fl;STOP-Tom pups or control STOP-Tom pups with low-titer serotype 8 adeno-associated virus (AAV8) that expressed Cre under the neuronal-specific Synapsin 1 promoter (AAV8-Syn-Cre; Fig. 6D). Serotype 8 AAV has been previously shown to have high tropism to Purkinje cells, allowing the cell type-specific expression of Cre and deletion of Amot in Purkinje cells (Gibson and Ma, 2011). Three weeks after the viral injection, we sacrificed the animals and analyzed the morphology of Purkinje cell dendritic trees using high-magnification images of 100 µm sagittal sections of the cerebellum (Fig. 6D, E). We measured the dendritic tree height and width, length of primary and secondary branches, and dendritic field area of Purkinje cells. To eliminate potential bias, all of the measurements were performed in a blinded manner and confirmed by two independent researchers. Notably, Amot^−/−^ Purkinje cells exhibited significant reductions of all of these parameters compared with control cells, indicating impairments in the dendritic arborization of Purkinje cells *in vivo* (Fig. 6E-I).

### Yap1 is required for Purkinje cell dendritogenesis

To investigate the function of Yap1 *in vivo* we crossed Yap1 fl/fl mice, in which exons 1 and 2 of Yap1 were flanked by loxP sites (Fig. 7A), with Syn-Cre transgenic mice. To verify exons excision, we performed genotyping of genomic DNA obtained from tail and brain. As shown in Fig. 7B, the electrophoresis of the genotyping reactions revealed a 697 bp PCR fragment that corresponded to the deleted Yap1 sequence for DNA that was obtained from the brain of Yap1 fl/fl;Syn-Cre (Fig. 7B). The Western blot analysis of cerebellar homogenates from homozygous mice expressing Cre additionally confirmed reduced level of Yap1 protein (Fig. 7C). Yap1 fl/fl;Syn-Cre mice were born according to the Mendelian ratio and exhibited no signs of lethality. Homozygous mice that expressed Cre, however, were significantly smaller and had a lower weight compared to their littermate controls of the same sex (Fig. 7D, E). Similarly to Amot mutants, Yap1 fl/fl;Syn-Cre mice appeared to have reduced cerebellar size (Fig. 7F) and cerebellar weight (Fig. 7G) compared with littermate controls.

**Figure 7.**
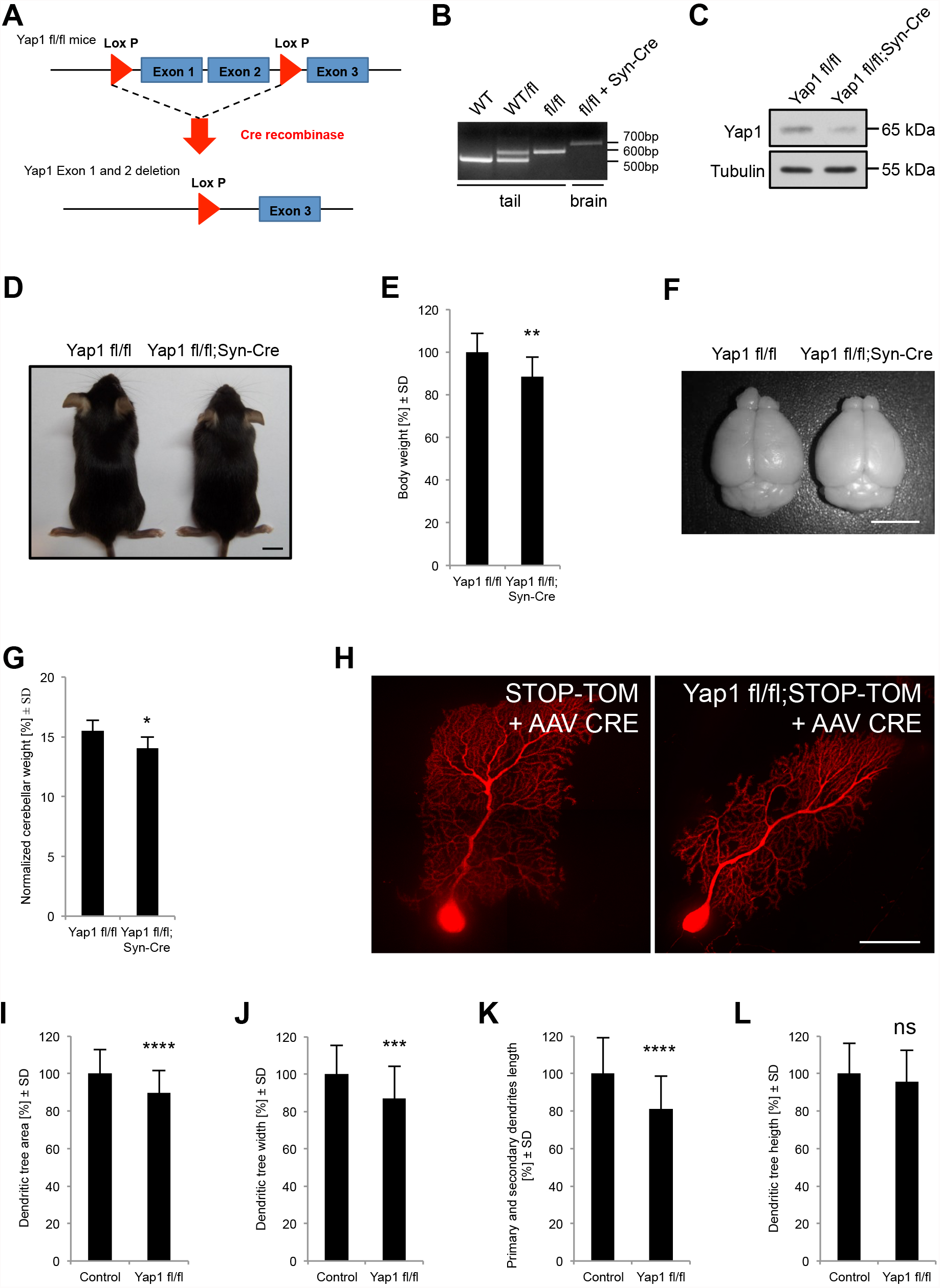
Yap1 deletion in neurons affects cerebellar size and impairs Purkinje cell dendritic tree morphology. (**A**) Strategy for generation of Yap1 conditional knockout mice. The Yap1 flox allele (upper diagram) and recombined allele after Cre-mediated excision (lower diagram). (**B**) Confirmation of Yap1 exon1-2 excision in the brain of Yap1 fl/fl;Syn-Cre mice assessed by PCR analysis of genomic DNA that was obtained from tail or brain. WT, wildtype control; WT/fl, Yap1 heterozygote; fl/fl Yap1homozygote mice. (**C**) Reduced Yap1 expression in the cerebellum of Yap1 fl/fl;Syn-Cre P30 mice analyzed by Western blot. (**D**) Yap1 fl/fl;Syn-Cre P30 mice appear smaller than Yap1 fl/fl control littermates; scale bars = 1 cm. (**E**) Weight analysis of Yap1 fl/fl (*n* = 20) and Yap1 fl/fl;Syn-Cre (*n* = 9) P30 mice; P = 0.0036. (**F**) Yap1 fl/fl;Syn-Cre mice exhibited a smaller cerebellum compared to control littermates (Yap1 fl/fl). Scale bars = 5 mm. (**G**) Quantitative analysis of cerebellum weight of Yap1 fl/fl (*n* = 10) and Yap1 fl/fl;Syn-Cre (*n* = 4) mice; P = 0.0178. Measurements were normalized to the whole brain weight. (**H**) Representative images of Purkinje cells from STOP-Tom (control) and Yap1 fl/fl;STOP-Tom mice infected with AAV8-Cre. Scale bar = 50 μm. Quantitative analysis of dendritic tree area (**I**), width (**J**), primary and secondary dendrite length (**K**) and height (**L**) of Purkinje cells from STOP-Tom (*n* = 33 cells and 5 mice) and Yap1 fl/fl;STOP-Tom (*n* = 80 cells and 8 mice) brains infected with AAV8-Cre; values shown as percentage of Control. P < 0.0001, P = 0.0003, P < 0.0001, P = 0.2194. Statistical significance was analyzed using two-tailed unpaired *t*-tests. ns, not significant; \**P* < 0.05, \*\**P* < 0.01, \*\*\**P* < 0.001, \*\*\*\**P* < 0.0001. Bars represent mean ± standard deviation (SD).

Finally, we investigated whether Yap1 also plays a role in the organization of Purkinje cell dendritic trees. We injected newborn Yap1 fl/fl;STOP-Tom mouse pups and control STOP-Tom mouse pups with the AAV8-Syn-Cre virus and analyzed Purkinje cell dendritic tree. To eliminate potential bias, all of the measurements were performed in a blinded manner and confirmed by two independent researchers. Yap1^−/−^ Purkinje cells exhibited a significant reduction of dendritic field area, dendritic tree width, and length of primary and secondary branches compared with control cells (Fig. 7H-K). We did not detected, however, significant differences in the dendritic tree height (Fig. 7L). Thus, Yap1, similar to Amot, plays an important role in regulating the dendritic arborization of Purkinje cells *in vivo*.

### Impaired motor coordination in mice with neuronal deletion of Amot or Yap1

To test whether morphological defects in Amot fl/fl;Syn-Cre or Yap1 fl/fl;Syn-Cre mice affect cerebellar function, we performed a motor coordination test using the rotarod. In this experiment, we placed animals on an accelerating rotarod and measured the latency to fall (Fig. 8A). To minimize differences in body size and weight in younger mice, we performed all of the tests on P100 animals. The latency to fall in mutant mice was significantly shorter in both Amot fl/fl;Syn-Cre and Yap1 fl/fl;Syn-Cre mice than in control animals of the same age and sex (Fig. 8B, C). This difference was apparent on the training day and during two consecutive days of the experiment. Thus, the smaller cerebellum of Amot fl/fl;Syn-Cre and Yap1 fl/fl;Syn-Cre mice and defects in Purkinje cell dendrite morphology in Amot−/− and Yap1−/− neurons were correlated with impairments in locomotor coordination.

**Figure 8.**
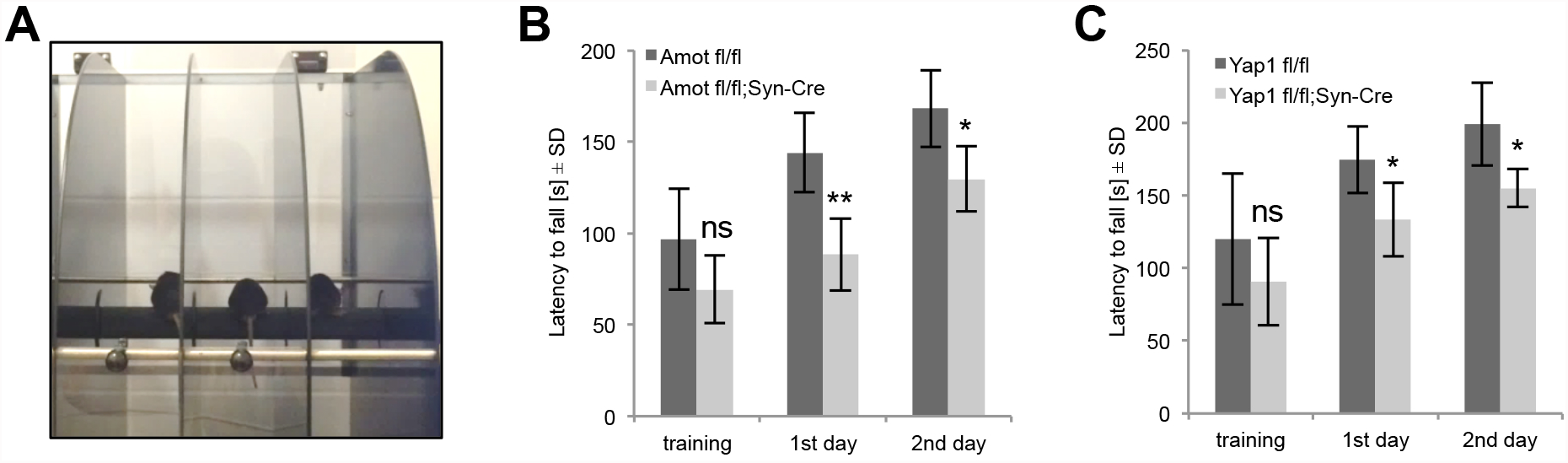
Impaired locomotor coordination in Amot fl/fl;Syn-Cre and Yap1fl/fl;Syn-Cre mice. (**A**) Image from the rotarod experiment. (**B**) Locomotor coordination of Amot fl/fl (*n* = 5) and Amot fl/fl;Syn-Cre (*n* = 5) P100 mice, revealed by the latency to fall from the rotarod; P = 0.0988, P = 0.0029 and P = 0.0137. (**C**) Locomotor coordination of Yap1 fl/fl (*n* = 5) and Yap1 fl/fl;Syn-Cre (*n* = 5) P80 mice, revealed by the latency to fall from the rotarod; P = 0.2611, P = 0.0270 and P = 0.0138. Statistical significance was analyzed using two-tailed unpaired *t*-tests. ns, not significant; \**P* < 0.05, \*\**P* < 0.01,. Error bars represent mean ± standard deviation (SD).

## DISCUSSION

It has been previously reported that Amotl2, one of the members of the Motin family of proteins, localizes to the postsynaptic machinery of the neuromuscular junction and regulates its organization (Proszynski and Sanes, 2013). Recently, Wigerius and colleagues (Wigerius et al., 2018) have reported that Amot localizes to the synapses in mature cultured neurons and regulates the postsynaptic specialization through the interaction with MUPP1 and PSD-95. In the present study we show that Amot distribution changes during neuronal development and at the early stages it localizes to dendrites and axons. Additionally, we show that Amot regulates dendritic tree morphogenesis in both cultured hippocampal cells and Purkinje neurons in the mouse cerebellum, what was associated with decreased cerebellar size. We show that one of known Amot-interacting proteins, transcription co-activator Yap1, is expressed in neurons and co-precipitates Amot from brain extract. Additionally, ectopic expression of Amot with mutated Yap1-binding sites fails to rescue phenotypes observed in Amot depleted neurons, suggesting that Amot regulates dendritic arborization through Yap1. Similarly to Amot, Yap1 localizes to dendritic processes in immature neurons (DIV 7) and relocalizes to synapses in fully developed cells (DIV 21) *in vitro* and adult brains. We have found that Yap1 knockdown or knockout, similar to Amot deletion, impairs the dendritogenesis of cultured hippocampal neurons, and the neuron-specific deletion of Yap1 in conditional knockout mice affects the organization of Purkinje cell dendrites and cerebellum size. Finally, neuronal deletion of either Amot or Yap1 leads to impaired locomotor coordination in rotarod test.

Although our results suggest that Amot regulates dendritic trees through Yap1, we cannot exclude that other mechanisms are also involved. Our data showed that transfection of neurons with truncated Amot constructs encoding either the ID domain alone, or Amot with deleted ID domain could partially rescue the defects in dendritic arbors caused by Amot depletion. These results suggest that different domains are important for Amot function in neurons. For example, CC domain plays a role in Amot oligomerization (Ernkvist et al., 2008; Wells et al., 2006), and PDZ-binding motif has been shown to recruit polarity proteins (e.g. Par3) (Ernkvist et al., 2009; Wells et al., 2006) that may also contribute to dendrite morphogenesis. On the other hand, the deletion of actin-binding region in Amot sequence did not attenuate its ability to rescue impairments in dendritic tree morphology in Amot knockdown neurons. This suggests that the interaction with actin is dispensable for Amot functions in dendritic development, but it could be critical for other Amot-mediated processes. For instance, Wigerius, et al. (Wigerius et al., 2018) have recently shown that in mature neurons Amot associates with actin and stabilizes actin cytoskeleton in dendritic spines, thus controlling the postsynaptic integrity and spines morphology. Therefore, depending on the developmental stage of neurons Amot could associate with different proteins and hence control different molecular processes.

Our analysis of mutant mice with neuronal deletion of Amot or Yap1 was concentrated on cerebellum because of its reduced size and the limitations of the Syn-Cre mice, in which Cre-mediated excision appeared to be the highest and the most homogeneous in cerebellar Purkinje cells. If broader Cre-mediated deletion could be achieved, the observed phenotypes in conditional knockout mice could be more pronounced. Interestingly, Huang and colleagues (Huang et al., 2016a) reported that Yap1 fl/fl mice that expressed Cre under the control of nestin regulatory elements (i.e., expression in progenitors of neural and glial cell lineages) had a substantially smaller brain size, but the authors interpreted this result as a consequence of the loss of Yap1 function solely in glial cells. Our localization as well as functional experiments including RNAi-mediated silencing and neuron-specific deletion of Yap1 using conditional-knockout mice suggest Yap1 expression and function in neurons. This is consistent with previous reports of Okazawa and coworkers (Mao et al., 2016; Yamanishi et al., 2017) showing that Yap1 is largely expressed in mouse and human neurons positive for Map2 and NeuN.

The best-characterized Yap1 function is being an effector protein in the Hippo signaling pathway that regulates cell-cell contact growth inhibition, mechanotransduction, adhesion, and cell proliferation including invasion of cancer cells. Little is known, however, about the function of Hippo signaling pathway in neurons. Emoto and coworkers (Emoto et al., 2006) suggested that Hpo and Wts (*Drosophila* orthologs of Mst1 and Lats1, two core components of the Hippo signaling pathway) play a major role in dendritic tree tiling and maintenance in *Drosophila* peripheral sensory neurons. Apart from Yap1, Amot also associates with other proteins that influence Hippo pathway signaling that could contribute to neuronal organization. For example, Merlin, which together with Amot forms a signaling platform that regulates phosphorylation of Mst and Lats was recently shown to play an important role in Purkinje cell dendritogenesis (Schulz et al., 2010).

In the Hippo signaling pathway Yap1 functions as one of downstream effector proteins. This transcriptional co-activator interacts with TEA-domain transcription factor (TEAD) and stimulates the expression of Hippo pathway-dependent genes (e.g., CTGF and Cyr61). In our study, we found that deletion of the TEAD binding region did not affect the ability of Yap1 to rescue the phenotype of Yap1 knockdown in neurons (Fig. 4c, d). Our observations suggest that the interaction between Yap1 and TEAD is dispensable for Yap1 function in dendrite morphogenesis, but we cannot exclude the possibility that the transcriptional activity of Yap1 is important for dendritic outgrowth or the regulation of other neuronal functions. Malik and coworkers (Malik et al., 2013) recently showed that Cyr61 is critical for the proper development of dendritic arbors in cultured hippocampal neurons. However, the authors did not link Cyr61 expression with the transcriptional activity of Yap1. Instead, they proposed a different mechanism, in which Cyr61 serves as a ligand for β1 integrin (Malik et al., 2013). Another possibility is that the interaction between Yap1 and TEAD in neurons has an important function only under certain physiological or pathological conditions. For example, Mao and colleagues (Mao et al., 2016) have proposed that Yap/TEAD-dependent necrosis is a major cause of neurodegenerative processes in Huntington’s disease. Thus, an interesting line of investigation would be to study the role of various Hippo pathway-related proteins in the context of the development and function of neuronal networks under various pathological conditions.

In addition to its role in Hippo pathway-dependent gene expression, Motins and Yap1 may also regulate other signaling pathways (Artinian et al., 2015; Azzolin et al., 2014; Huraskin et al., 2016; Imajo et al., 2012; Kim et al., 2017; Konsavage et al., 2012; Li et al., 2012; Tumaneng et al., 2012; Zhao et al., 2017). For example, Yap1, Amot, and two other Motins were shown to modulate the activity of Wnt/β-catenin signaling and regulate cell differentiation in tumor and healthy tissues (Azzolin et al., 2014; Huraskin et al., 2016; Imajo et al., 2012; Kim et al., 2017; Konsavage et al., 2012; Li et al., 2012; Zhao et al., 2017). Yap1 has also been shown to mediate crosstalk between the Hippo and PI(3)K-TOR signaling pathways that are involved in regulating organ size (Tumaneng et al., 2012). Recent studies by Artinian and coworkers (Artinian et al., 2015) showed that mechanistic target of rapamycin (mTOR) signaling might regulate Yap1 activity through Angiomotins. Elucidation of the exact context of these processes is of great interest to neuroscience and also to other disciplines of biology and medicine and will require additional extensive investigations in the future.

## MATERIALS AND METHODS

### DNA constructs and antibodies

β-actin-GFP plasmid (eGFP under the control of β-actin regulatory elements) was described previously (Jaworski et al., 2005). β-actin-mRFP plasmid (mRFP under the control of β-actin regulatory elements) was a gift from Jakub Włodarczyk (Nencki Institute of Experimental Biology, Warsaw, Poland) (Skupien et al., 2014). Amotp130-delAB was donated by Dannel McCollum (University of Massachusetts Medical School, Worcester, MA, USA) (Mana-Capelli et al., 2014). The following plasmids were obtained from Addgene (Cambridge, MA, USA): pLKO.1 (catalog no. 10878)(Moffat et al., 2006), pEGFP-C3-hYAP1 (catalog no. 17843) (Basu et al., 2003), pLX304-YAP1_60-89 (catalog no. 59144) (Shao et al., 2014), HA-AMOT p130 Y242/287A (Amot ΔYap1, catalog no. 32822) (Zhao et al., 2011). pCMV-Flag-Amot was a gift from Chunling Yi (Georgetown University Medical Center, Washington, DC, USA) (Yi et al., 2013). Yap1, and Amot constructs were PCR-amplified and cloned into the β-actin-GFP vector to generate GFP fusion proteins. The pFUCherry-TA(3x)-iCre plasmid was a gift from Julie Lefebvre (The Hospital for Sick Children, University of Toronto, Toronto, ON, Canada). The shRNA sequences 5’-GGTGGAGACTGAAATCCAACG-3’ (Amot shRNA) and 5’-GAAACAGCAGGAGTTATTTCG-3’ (Yap1 shRNA) were cloned into the pLKO.1 vector.

The following antibodies were used: anti-α tubulin (Abcam, Cambridge, MA, USA; catalog no. ab18251), anti-Yap1 (Sigma-Aldrich, St. Louis, MO, USA; catalog no. WH0010413M1), anti-Amot (Aviva, San Diego, CA, USA; catalog no. ARP34661_P050), anti-Amot (Santa Cruz Biotechnology, Dallas, TX, USA; catalog no. sc-515262) anti-Map2 (Sigma, St. Louis, MO, USA; catalog no. M1406), anti-Yap1 (Developmental Studies Hybridoma Bank, Iowa City, IA, USA; catalog no. 8J19), anti-Synaptophysin 1 (Synaptic Systems, Goettingen, Germany; catalog no. 101004), and anti-Calbindin (Synaptic Systems, Goettingen, Germany; catalog no. 214006). Anti-Amot antibody was a gift from Lars Holmgren (Karolinska Institutet, Stockholm, Sweden) (Ernkvist et al., 2006). Horseradish peroxidase-conjugated anti-mouse immunoglobulin G (IgG) and anti-rabbit IgG were obtained from Cell Signaling (Danvers, MA, USA; catalog no. 7076 and 7074, respectively). Alexa-Fluor-488-, Alexa-Fluor-568-, and Alexa-Fluor-647-conjugated secondary antibodies were obtained from Thermo Fisher Scientific (Waltham, MA, USA; catalog no. A-11034, A-11004, A-11041, and A-21450). DAPI was purchased from AppliChem (Darmstadt, Germany; catalog no. A4099).

### Cell culture and transfection

Primary hippocampal and cortical neuronal cultures were prepared from Wistar rat and C57BL/6 mouse brains that were collected on embryonic day 18 (E18). The cultures were performed according to the procedure of Banker and Goslin (Banker and Goslin, 1988). Neurons were dissociated with trypsin and cultured in Neurobasal medium (Thermo Fisher Scientific, Waltham, MA, USA; catalog no. 21103049) supplemented with 2% B27 (Thermo Fisher Scientific, Waltham, MA, USA; catalog no. 17504044), L-glutamine, and glutamic acid and plated on poly-D-lysine (Sigma-Aldrich, St. Louis, MO, USA; catalog no. P2636) and laminin (Invitrogen, Grand Island, NY, USA; catalog no. 23017-015) coated surfaces. Primary hippocampal neurons were transfected with Lipofectamine 2000 (Thermo Fisher Scientific, Waltham, MA, USA; catalog no. 11668-030) at 7 days *in vitro* (DIV7) as described previously (Perycz et al., 2011). Amaxa nucleofection was performed with an Amaxa Nucleofector II Device (Lonza Cologne AG, Cologne, Germany; catalog no. AAD-1001S) using the Amaxa Rat Neuron Nucleofector Kit (Lonza Cologne AG, Cologne, Germany; catalog no. VPG-1003) according to the manufacturer’s protocol.

### RNA isolation and quantitative real-time PCR

RNA was isolated using TRIsure reagent (Bioline, Taunton, MA, USA; catalog no. BIO-38033), and cDNA was synthesized using a High-Capacity cDNA Reverse Transcription Kit (Thermo Fisher Scientific, Waltham, MA, USA; catalog no. 4368814) according to the manufacturer’s instructions. qRT-PCR was performed using the Step One Plus real-time PCR system (Thermo Fisher Scientific, Waltham, MA, USA) and SYBR Green PCR Master Mix (Thermo Fisher Scientific, Waltham, MA, USA; catalog no. 4309155). Manufacturer programs were used for data acquisition, and the results were analyzed by the comparative *Ct* method for relative quantification. The housekeeping gene Glyceraldehyde-3-phosphate dehydrogenase (Gapdh) was used to standardize the samples. The following primers were used: Gapdh (5’-GGCCTTCCGTGTTCCTAC-3’ and 5’-TGTCATCATACTTGGCAGGTT-3’), Amot (5’-CAGTCATTAGCCACTCTCCTAAC-3’ and 5’-GTCTTAATCCTTCCTTCCATGTC-3’).

### SDS-PAGE and Western blot

Cultured cells or brain tissues were lysed with radioimmunoprecipitation assay (RIPA) buffer (50 mM Tris-HCl [pH 8.0], 150 mM NaCl, 2% Igepal CA-630 [NP-40], 0.25% sodium deoxycholate, 1 mM NaF, and 1 mM DTT supplemented with Mini Protease Inhibitors cocktail [Roche, Indianapolis, IN, USA; catalog no. 11873580001]). Proteins were separated by sodium dodecyl sulfate-polyacrylamide gel electrophoresis (SDS-PAGE) and transferred to nitrocellulose membranes (PALL, Port Washington, NY, USA; catalog no. 66485) using the Trans-Blot Turbo system (Bio-Rad, Hercules, CA, USA; catalog no. 1704270). After blocking with 5% milk in Tris-buffered saline with Tween (TBST), primary antibodies were added to the blocking buffer. The membranes were then washed with TBST and incubated with the appropriate horseradish peroxidase-conjugated secondary antibodies.

### Co-immunoprecipitation

Mouse brain tissue was homogenized in ice-cold lysis buffer (50 mM Tris-HCL [pH 8], 150 mM NaCl, 0,5% Igepal CA-630 [NP-40], 10% glycerol, 1 mM DTT, 1 mM NaF, and Mini Protease Inhibitor cocktail [Roche, Indianapolis, IN, USA; catalog no. 11873580001]). The lysate was then passed three times through a 25-gauge needle with a syringe, rotated at 4°C for 15 min, and centrifuged for 30 min at 18,000 × g at 4°C. Supernatant from centrifugation was incubated overnight with Dyneabeads Protein G (Thermo Fisher Scientific, Waltham, MA, USA; catalog no. 10004D) coated with anti–Yap1 antibody (Sigma-Aldrich, St. Louis, MO, USA; catalog no. WH0010413M1). Beads with attached proteins were washed four times with lysis buffer, resuspended in 2×sample buffer and boiled for 10 minutes. For Western blot analysis, samples were subjected to the SDS-PAGE electrophoresis.

### Immunocytochemistry and immunohistochemistry

Hippocampal neurons were grown on glass coverslips, fixed with 4% paraformaldehyde (PFA), and blocked with blocking buffer (2% bovine serum albumin [BSA], 2% normal goat serum, and 0.5% Triton X-100 in PBS). The primary and secondary antibodies were applied in blocking buffer.

For brain immunohistochemistry, the mice were transcardially perfused with ice-cold PBS, followed by perfusion with 4% PFA in PBS. The brains were removed, post-fixed in 4% PFA overnight at 4°C, and cryoprotected in 30% sucrose. Coronal brain slices (40 µm) were obtained with a cryostat (Leica, Wetzlar, Germany; Leica 1950 Ag Protect) and collected in antifreeze solution (30% ethylene glycol, 15% sucrose, and 0.05% sodium azide in PBS). After incubation in blocking buffer (10% BSA and 0.3% Triton-X in PBS), primary antibodies that were diluted in blocking buffer were applied. Fluorophore-conjugated secondary antibodies were diluted in 0.3% Triton-X in PBS.

### Mouse strains and husbandry

All of the mice were on a C57BL/6 genetic background and bred in a specific pathogen-free (SPF) animal facility at the Nencki Institute of Experimental Biology. Amot fl/fl;Syn-Cre and Yap1 fl/fl,Syn-Cre mice were generated by crossing Amot fl/fl (Yi et al., 2013) and Yap1 fl/fl mice (Zhang et al., 2010), respectively, with Synapsin-1 Cre (Syn-CRE) transgenic mice (B6.Cg-Tg[Syn1-cre]671Jxm/J). Syn-CRE and Yap1 fl/fl mice (*Yap1^tm1.1Dupa^*/J) were obtained from Jackson Laboratory (Bar Harbor, ME, USA; catalog no. 003966 and 027929, respectively). For single Purkinje cell analysis, Amot fl/fl and Yap fl/fl mice were crossed with the STOP-tdTomato (STOP-Tom) reporter strain (Madisen et al., 2010). All of the experimental procedures were approved by the First Warsaw Local Ethics Committee for Animal Experimentation, and the experiments were conducted in accordance with all applicable laws and regulations. The following primers were used for genotyping: Amot (5’-GATGGATGCTATGAGAAGGTG-3’ and 5’-GTAAGGATTACAGAGTCTGGG-3’; wildtype, ∼400 bp; flox, ∼500 bp; conditional knockout, no fragments), Amot (5’-ATAGCTAGTGAGCAGTAGCAG-3’ and 5’-GTAAGGATTACAGAGTCTGGG-3’; wildtype, ∼1 kbp; flox, ∼1.2 kbp; conditional knockout, ∼400 bp), and Yap1 (5’-CCATTTGTCCTCATCTCTTACTAAC -3’ and 5’-GATTGGGCACTGTCAATTAATGGGCTT-3’ and 5’-CAGTCTGTAACAACCAGTCAGGGATAC-3’; wildtype, ∼498 bp; flox, ∼597 bp; conditional knockout, ∼697 bp).

### Viral infection

Newborn pups (P0) were intracerebrally injected with 1.7 × 10^9^ viral genomes (VG) of AAV8-Syn-Cre (Signagen Laboratories, Rockville, MD, USA; catalog no. SL100883) as described previously (Gibson and Ma, 2011). Three weeks after viral infection, the animals were intracardially perfused with 4% PFA as described above, and brains were harvested and sectioned into 100 µm coronal slices.

### Rotarod test

To assess motor coordination, a computer-interfaced rotarod that accelerated from 4 to 40 rotations per min over 240 sec was used (TSE, Bad Homburg, Germany). The shaft diameter was 30 mm. The animals were taken to the behavioral testing room 1 h prior to the measurements on consecutive days to allow habituation, and habituation began 1 day before beginning the experiment. The animals were allowed to stand on the rotarod for 10-20 s before it began to rotate. The mice were tested in three trials per day, with a 20-min intertrial interval, over 3 consecutive days. The time that each mouse maintained its balance on the rotating rod was measured as the latency to fall.

### Confocal microscopy, image analysis, and quantification

Fluorescence microscopy was performed at the Confocal Microscopy Facility, Nencki Institute, using a spinning disc confocal microscope (Zeiss, Jena, Germany) equipped with PLN APO 10×/0.45, C APO 40×/1.20, and PLN APO 63×/1.40 objectives or Zeiss LSM800 Airyscan (Zeiss, Jena, Germany) equipped with PLN APO 63x/1.40 objective and Airyscan Detector (32x GaAsP detectors). Images were collected using ZEN software (Zeiss, Jena, Germany) and analyzed using ImageJ software (Schneider et al., 2012).

The morphometric analysis and quantification of dendritic tree complexity were performed using ImageJ software with the Neuronal Tracer plug-in to calculate the TDL and Sholl plug-in for Sholl analysis. Briefly, each image was processed with the Neuronal Tracer plug-in, and a mask of all dendrites on a confocal image was drawn manually. To avoid background that could be present in the original pictures, the mask was saved as a separate image and used in the automated Sholl analysis. For Purkinje cell analysis, confocal Z-stacks were collected and orthogonal projections were generated. Purkinje cell dendritic arbor area, width, height, and length were manually traced using ImageJ software. To eliminate potential bias, all of the measurements were performed in a blinded manner and confirmed by two independent researchers.

### Statistical analysis

Quantitative data are presented as the mean ± SEM or SD. P values and statistical methods (two-tailed and unpaired t-test, two-way ANOVA with Bonferroni’s post hoc test) are defined in the figure legends. All of the observations and analyses were performed based on at least three independent experiments. Quantitative analysis of Purkinje cell morphology, rotarod test and cerebellum weight analysis were performed in a blinded manner and confirmed by two independent researchers. Animals were assigned to experimental groups based on their genotypes. The statistical analyses were performed using GraphPad Prism 7 and Microsoft Excel software.

The data that support the findings of this study are available from the corresponding author upon reasonable request.

## ACKNOWLEDGEMENTS

Preparation of this article was supported by a Sonata-Bis grant (no. 2012/05/E/NZ3/00487 awarded to T.J.P.) and Preludium grant (no. 2015/19/N/NZ3/02346 awarded to K.R.) from the Polish National Science Center (NCN). L.H. is supported by grants from the Swedish Cancer Society, Swedish Research Council and the Cancer Society of Stockholm.

## AUTHORS’ CONTRIBUTIONS

K.R conceived and performed all of the cell cultures and molecular experiments, analyzed the data, and wrote the manuscript. J.K. conceived and performed all of the *in vivo* experiments, including the behavioral tests and virus experiments, analyzed the data, and contributed to manuscript preparation. H.D. and J.J provided initial help with the neuron cultures and Amaxa experiments. W.K. provided STOP-Tom mice. L.H. provided Amot conditional knockout mice. P.B., L.K., W.K., M.R., J.J., and L.H helped design the experiments and analyze the data and provided comments on the manuscript. T.P. coordinated the project, designed the experiments, analyzed the data, and contributed to writing the manuscript and preparing the figures. We thank Jaworski’s Lab members for their help with experiments on cultured neurons.

